# Human urine alters methicillin-resistant *Staphylococcus aureus* virulence and transcriptome

**DOI:** 10.1101/2021.01.14.426765

**Authors:** Santosh Paudel, Kamal Bagale, Swapnil Patel, Nicholas J. Kooyers, Ritwij Kulkarni

## Abstract

Gram-positive methicillin-resistant *Staphylococcus aureus* (MRSA) is an emerging cause of hospital-associated urinary tract infections, especially in catheterized individuals. Despite being rare, MRSA-UTI are prone to potentially life-threatening exacerbations such as bacteremia that can be refractory to routine antibiotic therapy. Hence, MRSA-UTI is an important of research topic. To delineate molecular mechanisms governing MRSA urinary pathogenesis, we exposed three *S. aureus* strains, of which two were MRSA, to human urine and analyzed virulence characteristics and gene expression. We also analyzed MRSA-1369 transcriptome following cultivation in human urine for 2h. Our results reveal that human urine induces global changes in MRSA transcriptome, marked by changes in genes encoding proteins involved in metabolic pathways, virulence, and transcriptional regulators. In addition, *in vitro* assays also showed that human urine alters, in a strain-specific manner, adherence to human bladder epithelial cells and fibronectin, hemolysis of sheep RBCs, and surface hydrophobicity. In summary, our results provide first important insights into how the urine may specifically alter MRSA physiology in turn facilitating MRSA survival in the nutrient-limiting and hostile urinary microenvironment.

**Importance:** Methicillin-resistant *Staphylococcus aureus* (MRSA) is an uncommon cause of urinary tract infections (UTI) in the general population. However, it is important to understand MRSA pathophysiology in the urinary tract because isolation of MRSA in urine samples is often secondary to potentially life-threatening MRSA bacteremia. In this report, we describe that cultivation in human urine alters MRSA global gene expression and virulence. We hypothesize that these alterations may aid MRSA adapt to the nutrient-limiting, immunologically hostile conditions within the urinary tract.

## Introduction

Gram-positive pathogen *Staphylococcus aureus* is an emerging cause of urinary tract infections (UTI) representing 1% of uncomplicated cases and 3% of complicated cases associated with the physical obstruction of the urinary tract [1]. Urinary catheterization is the single most important predisposing factor for persistent *S. aureus* colonization of the urinary tract, which in turn substantially increases the risk of symptomatic urinary tract infection and potentially life-threatening, invasive sequelae such as bacteremia, endocarditis and septic shock [2, 3]. In addition, up to 20% of *S. aureus* catheter-associated UTI (CAUTI) are caused by methicillin-resistant *S. aureus* (MRSA), which are resistant to routine antibiotic therapy [4]. Hence, defining the complex interactions between host immune defenses and pathogen virulence effectors during the course of MRSA UTI is a clinically relevant research area despite relatively low prevalence of MRSA UTI.

Previous experiments have identified bacterial virulence factors central to the urinary pathogenesis of *S. aureus*. Examination of *S. aureus* isolated from catheterized patients has revealed that ∼80% of the clinical isolates form biofilm; a virulence characteristic apparently associated with the presence of *icaA* and *icaD* genes encoding polysaccharide capsule [5]. The trace metal nickel/cobalt transporter systems are implicated in the urinary fitness and virulence of *S. aureus* due to the involvement of nickel as a cofactor in urease enzyme activity [6, 7]. MRSA infection is shown to exacerbate catheterization-induced bladder inflammation in a mouse model by inducing prolonged production of potent pro-inflammatory cytokines (IL-1α, IL-1β, IL-6, IL-17, and TNFα) and the recruitment of macrophages and neutrophils [8]. In addition, MRSA also facilitates bladder colonization by inducing accumulation of host protein fibrinogen on urinary catheters, to which it adheres via surface adhesins clumping factor A and B (clfA, clfB) [8]. Collectively, these studies have identified virulence factors that afford survival advantage to MRSA inside the urinary tract and immune responses that defend the host. However, these studies were not designed to define the pleiotropic effects of urinary microenvironment on MRSA physiology, which is the main objective of our project as reported here.

For delineating complex mechanisms regulating the survival of uropathogens in the human urinary tract and their ability to cause UTI, the ideal experimental setup would be to examine bacterial transcriptome and proteome in real-time through different stages of UTI in a human host. Given the obvious impracticability of this setup, in this report we explored a more practical alternative and analyzed MRSA physiology in human urine *in vitro*. We used three *S. aureus* strains, namely MRSA-1369 and PUTS-1, which are clinical isolates from urine [8], and USA300, which has emerged in the last two decades as the predominant community-associated MRSA strain in the US [9]. We compared these strains cultivated in human urine with control in nutrient rich culture medium by *in vitro* virulence assays. In addition, we used RNA sequencing (RNASeq) to compare global transcriptomic profiles of MRSA-1369 cultivated in human urine or in nutrient-rich tryptic soy broth (TSB). Our results reveal that in addition to myriad metabolic adaptations necessary for survival in nutrient-limiting conditions in urine, MRSA cultivated in human urine also exhibits induction of virulence characteristics such as adherence to uroepithelial cells, hydrophobicity, and hemolysis. Overall, our study provides important, first insights into the effects of urinary microenvironment on MRSA physiology.

## Results

### *S. aureus* strains grow in human urine

We compared CFUs in TSB control and human urine at different time points up to 24h to confirm that MRSA-1369, PUTS-1, and USA300, were not inhibited by human urine (Fig 1). However, the doubling times of all three strains cultivated in human urine exhibited 1.4 to 2.4-fold reduction (*P*<0.05, unpaired T test) in comparison to the doubling times of TSB-controls (Fig 1). Also, the doubling time for PUTS-1 (87±19 min) in human urine was higher than the doubling time for MRSA-1369 (137±37 min; *P*=0.1) and USA300 (156±7 min, *P*<0.05).

**Figure 1.**
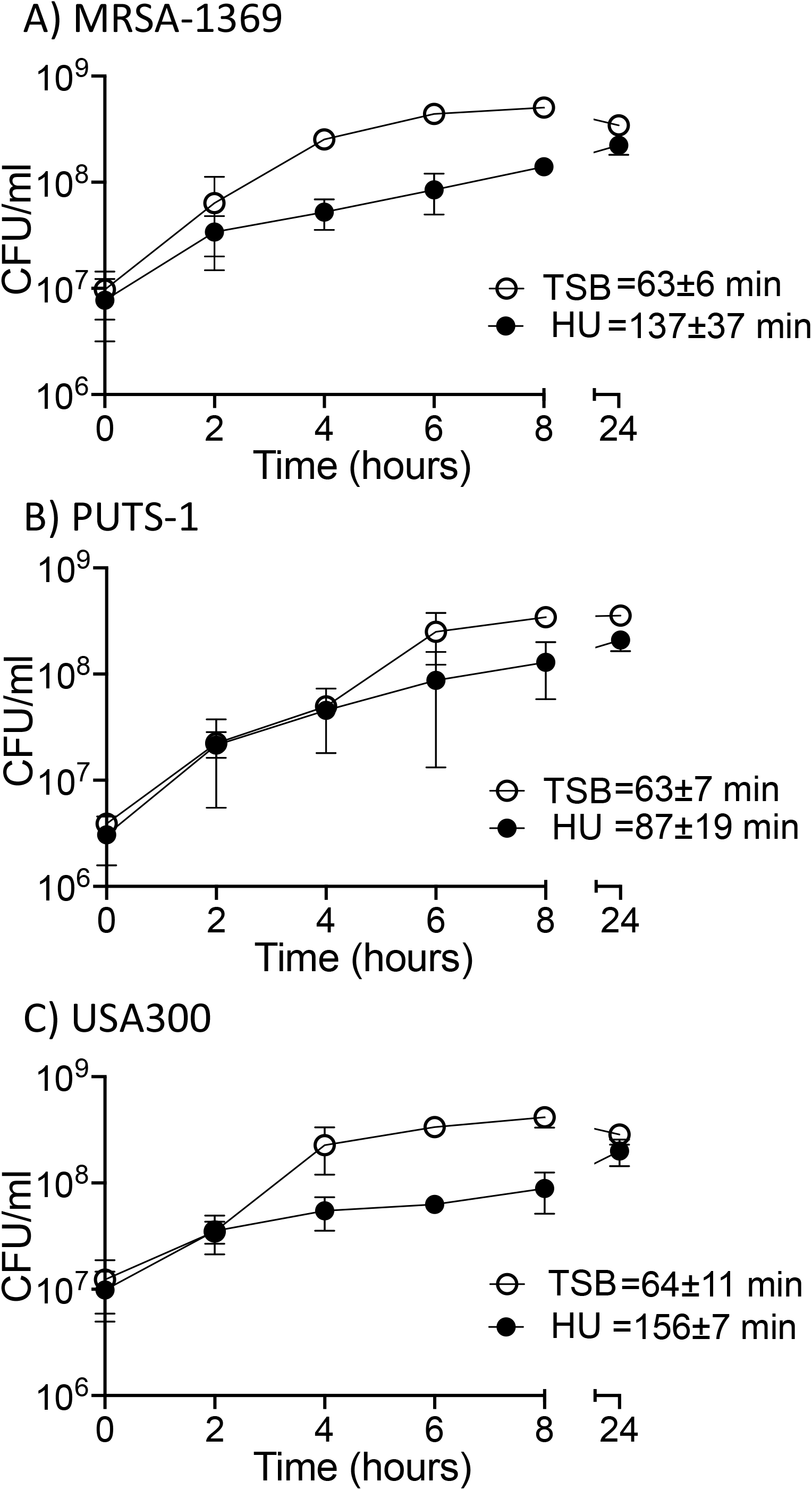
*S. aureus* strains are able to grow in human urine. (A) MRSA-1369, (B) PUTS-1, and (C) USA300 were cultivated either in TS broth (TSB) or human urine (HU). Bacterial growth was monitored by CFU enumeration over a period of 24 hours. For each time point, average CFU/ml (three biological replicates) ± StDev is shown. Also shown is average doubling time ± StDev for each strain in TSB or in human urine.

### Effects of human urine on the virulence characteristics of *S. aureus* strains

MRSA produces a large arsenal of virulence factors to facilitate its colonization, persistence, and dissemination within the host and for evasion of immune defenses. Hence, in the next set of experiments, we examined the effects of 2h-long exposure to human urine on the virulence characteristics of *S. aureus* strains.

Bacterial adherence is the first crucial step in an infection as it facilitates colonization and subsequent invasion across the mucosal barrier. We compared *S. aureus* strains cultivated in human urine with the control for the ability to adhere to 5637 human bladder epithelial cell line and to extracellular matrix protein fibronectin (Fig 2). Human urine significantly increased adherence of MRSA strains MRSA-1369 and USA300 to bladder epithelial cells (Fig 2A, C). In contrast, % adherence was similar for PUTS-1 irrespective of whether it was cultivated in human urine or TSB control (Fig 2B). When cultivated in human urine for 2h, all three strains showed decreased binding to fibronectin, however this change was statistically significant only for USA300 (Fig 2 D, E, F).

**Figure 2.**
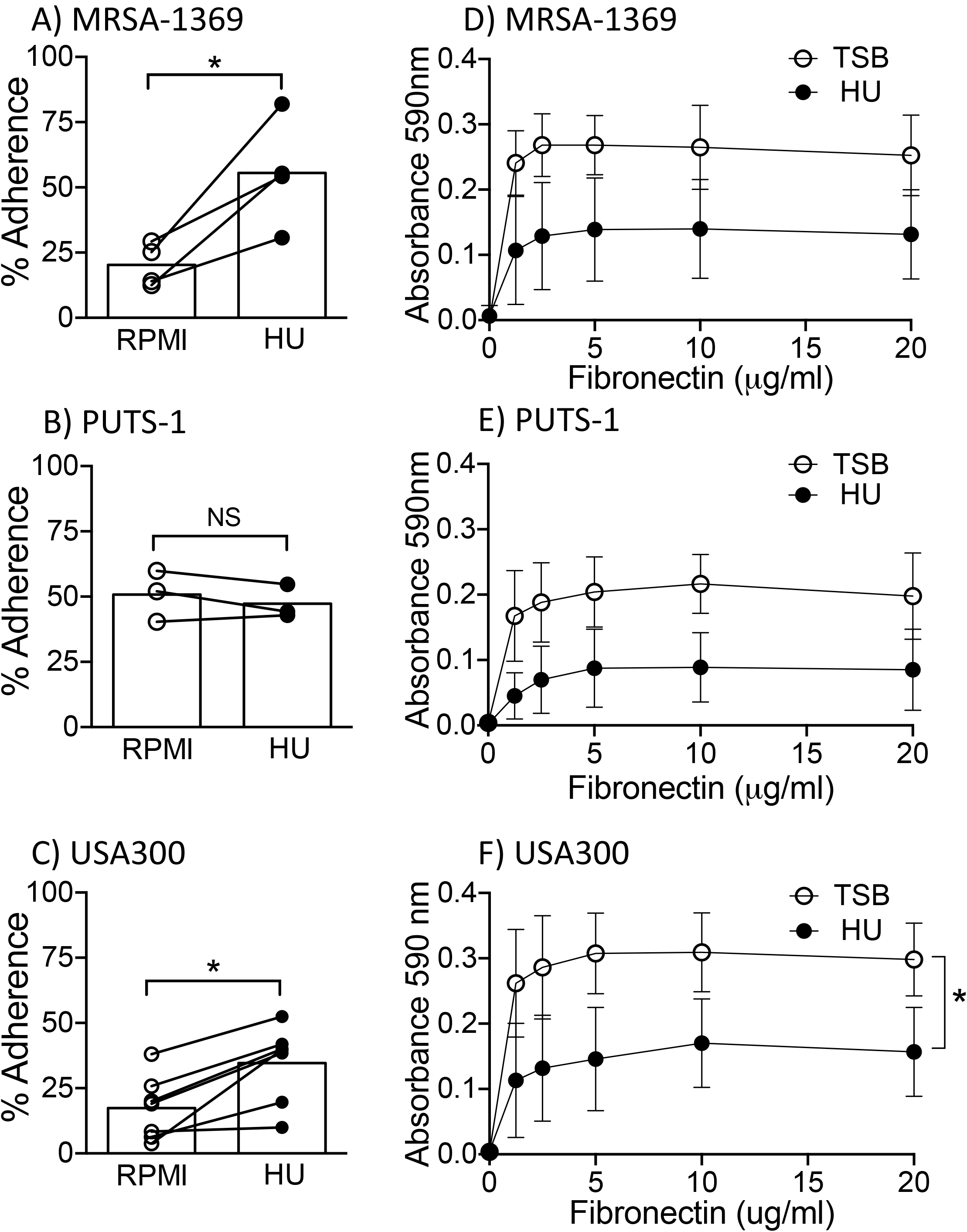
Human urine alters *S. aureus* adherence to biotic material in a strain-dependent manner. (A) MRSA-1369, (B) PUTS-1, and (C) USA300 were added to the monolayers of 5637 human bladder epithelial cell line either in the presence of plain RPMI (control) or human urine (HU). After 2h-long incubation at 37°C, cell adherent bacteria were enumerated. % adherence for biological replicates (each with two or more technical replicates) are reported as scatter diagram with mean shown as histogram. Statistical significance was determined by paired t test. In separate experiments, (D) MRSA-1369, (E) PUTS-1, and (F) USA300 pre-exposed to either TSB or HU for 2h were incubated in plates coated with human fibronectin. After vigorous washing, bacteria adherent to fibronectin were quantified by crystal violet staining and measurement of absorbance at 590nm. Mean absorbance ± StDev values are reported for each fibronectin concentration and compared by paired t test. For all figures * refers to *P*<0.05.

Next, we used microbial adhesion to hydrocarbons test (MATH) [10] to analyze changes in staphylococcal surface hydrophobicity caused by human urine. In this assay, TSB-control and human-urine-pre-exposed *S. aureus* strains were mixed with a hydrocarbon, hexadecane. At the end of 30 min incubation time, bacteria present in the aqueous phase were enumerated to estimate hydrophobicity. In comparison to control, significantly lower numbers of human urine exposed MRSA-1369 were found in the aqueous phase indicating that human urine increases hydrophobicity of MRSA-1369 (Fig 3A). Reduction in surface hydrophobicity is an important immune evasion mechanism used by staphylococci to avoid killing by antimicrobial peptides [11]; however, we did not observe corresponding changes in the killing of MRSA-1369 by human cathelicidin, LL37 (data not shown). Exposure to human urine did not alter surface hydrophobicity of either PUTS-1 (Fig 3B) or USA300 (Fig 3C).

**Figure 3.**
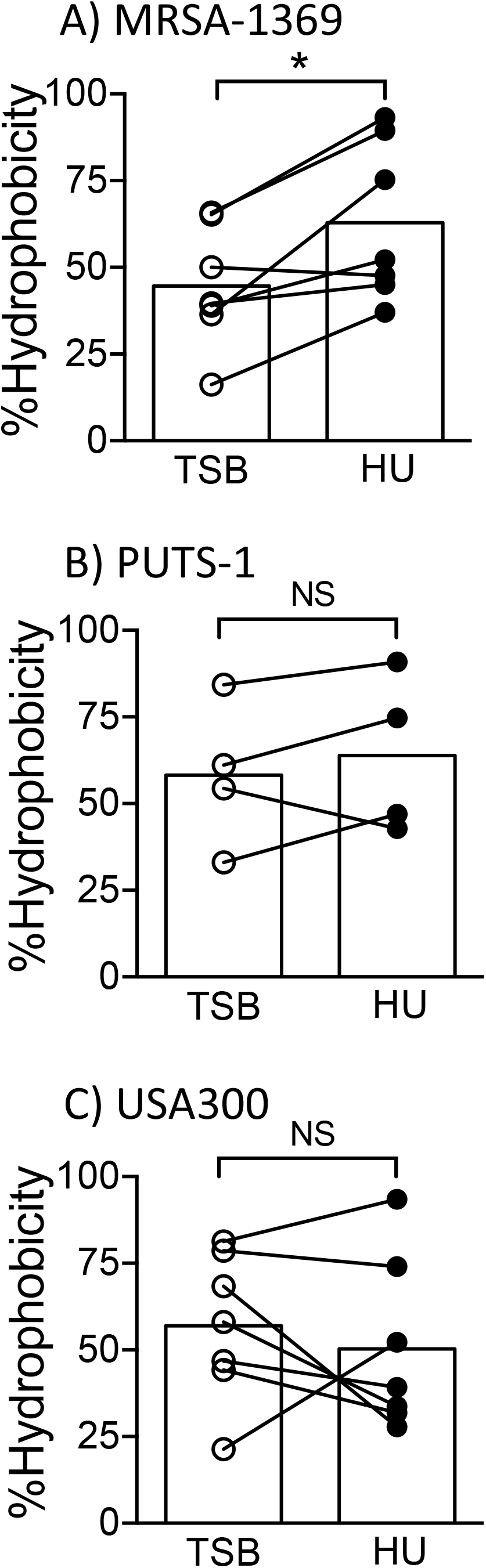
Effects of human urine on *S. aureus* hydrophobicity. Following pre-exposure to either TSB or human urine (HU), surface hydrophobicity was quantified for (A) MRSA-1369, (B) PUTS-1, and (C) USA300. % hydrophobicity (with average shown as histogram) are reported. Statistical significance was determined by paired t test.

*S. aureus* secretes numerous cytolytic toxins to target and kill mammalian cells by damaging their plasma membrane. The cytolytic activity contributes to pathogenesis by helping MRSA evade phagocyte-mediated killing [12]. To visualize whether exposure to human urine alters staphylococcal cytolytic activity, we incubated two-fold dilutions of *S. aureus* strains (starting from 10^8^ CFU/ml) pre-exposed for 2h to either human urine or TSB-control with sheep RBCs at 37°C. After incubation, we quantified hemoglobin released in the supernatant spectrophotometrically. Both MRSA-1369 and USA300 pre-exposed to human urine exhibited ∼1.5-fold higher hemolysis compared to TSB controls (Fig 4). In a striking contrast, control PUTS-1 exhibited very low hemolysis activity, which was not altered by pre-exposure to human urine.

**Figure 4.**
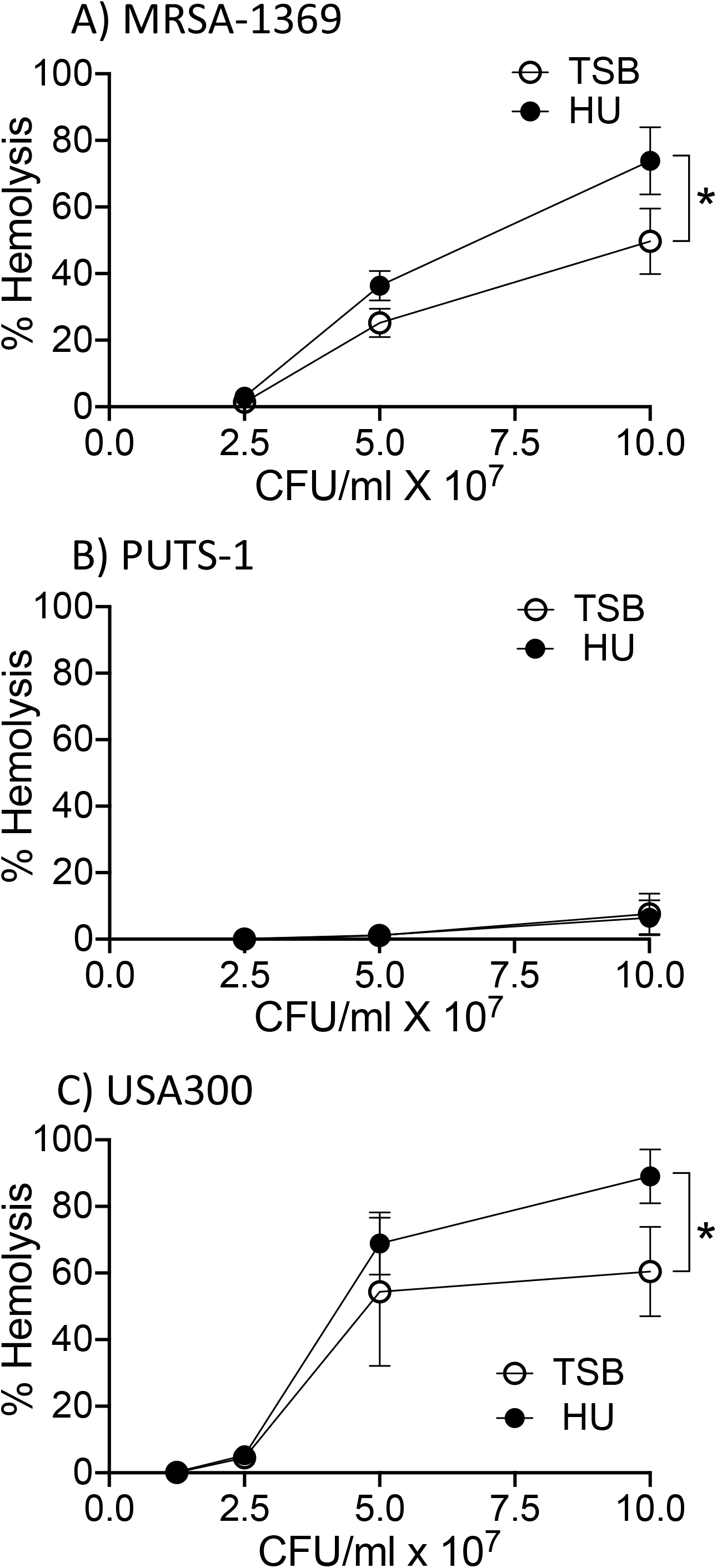
Human urine induces lysis of sheep RBCs by *S. aureus* in a strain-dependent manner. Mid-log cultures of (A) MRSA-1369, (B) PUTS-1, and (C) USA300 were cultivated for 2h in either TS broth (TSB) or in human urine (HU). Two-fold dilutions of bacteria (from 10^8^ CFU/ml to 1.25×10^7^ CFU/ml) were then incubated with sheep RBCs for 2h at 37°C. Intact RBCs were centrifuged. The absorbance at 420nm of supernatant was reported as % of total hemolysis (induced by tritonX100 treatment). Average % hemolysis (at least three biological replicates, each with at least two technical replicates) ± StDev is reported. Statistical significance was determined by paired t test.

Staphylococcal cell wall homeostasis involves two competing processes, namely cell wall synthesis governed by cell wall synthesizing enzymes (*aka* penicillin binding proteins) and autolysis regulated by murein hydrolases (*aka* autolysins). *S. aureus* autolysis is upregulated in response to adverse physiological conditions including exposure to cell wall inhibitor β lactam antibiotics [13]. We did not observe significantly changes in autolysis in any of the staphylococcal strains following 2h-long cultivation in human urine (data not shown).

### Effects of human urine on the gene expression in *S. aureus* strains

Using quantitative real-time PCR, we compared expression of select genes from MRSA-1369, PUTS-1, and USA300 following exposure to human urine for 2h (Fig 5). The genes selected for this analysis include virulence genes *atl* (autolysin), *clfAB* (clumping factor A and B), *fib* (fibrinogen binding protein), *fnbA* (fibronectin binding protein A), and *hla* (hemolysin) and transcriptional regulators *agrC* (accessory gene regulator) and *sarA* (staphylococcal accessory regulator). Genes that were two-fold up— or down—regulated were considered significant. As shown in Fig 5 human urine altered in a strain-specific manner the expression of *agrC, clfA, fib* (≥ 2-fold altered in PUTS-1, unchanged in others), and *hla* (≥ 3-fold upregulated in MRSA-1369 and USA300, 2.3-fold upregulated in PUTS-1). In contrast, the expression of *clfB* and *fnbA* was downregulated by human urine to a similar extent in all three *S. aureus* strains.

**Figure 5.**
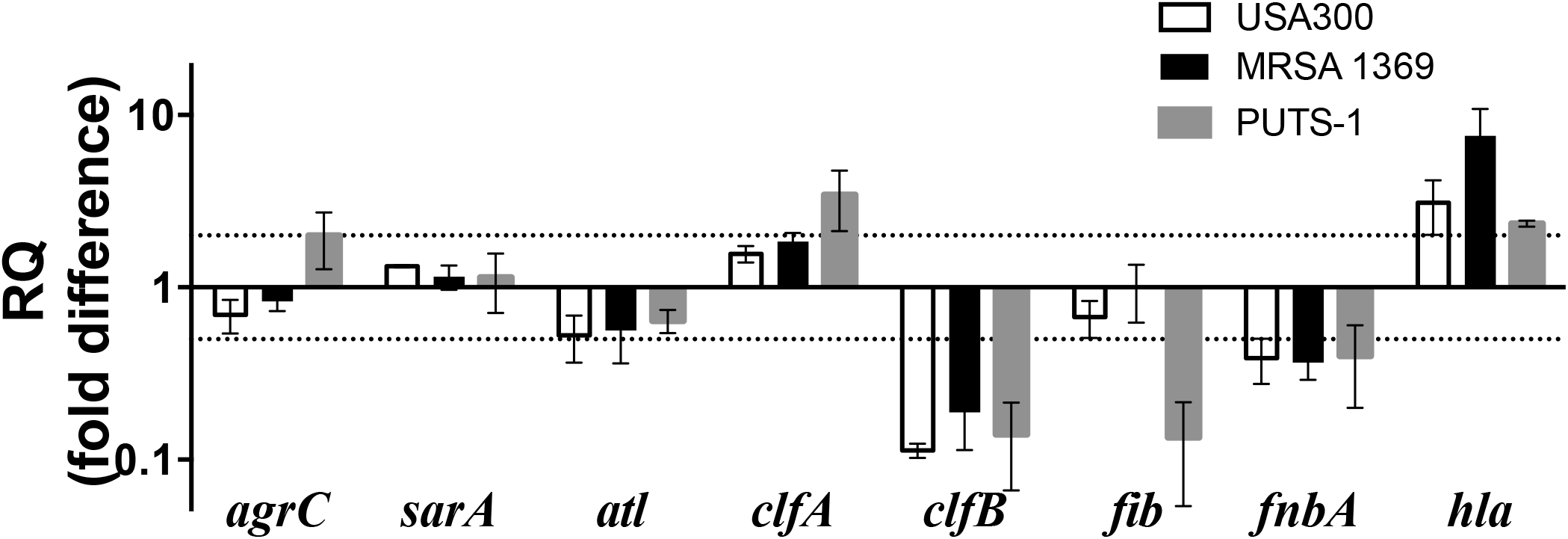
qRTPCR results for *S. aureus* strains following cultivation in human urine. Transcript levels for specific virulence and associated genes (indicated on *x*-axis) in MRSA-1369, PUTS-1, and USA300 were determined by quantitative real time PCR with normalization to 16SrRNA. RQ values were calculated by comparative threshold cycle (ΔΔC_T_) algorithm. RQ fold differences over transcript levels from TSB-control are presented as average of from at least two biological replicates (each with three technical repeats)± standard deviation are shown. Dotted lines indicate 2 fold up—or down—regulation.

### RNA-Seq and read mapping

We used RNAseq to quantify differential gene expression by assessing variation across the transcriptome for MRSA-1369 cultivated in human urine vs. in TSB. MRSA-1369 was selected for RNASeq analysis, as it is a clinical isolate from a patient suffering from CAUTI [8]. RNA was isolated from three independent biological replicates in each treatment. Whole transcriptome sequencing with rRNA depletion resulted in an average of 25.2 million 150bp paired-end reads per sample (range: 21.6-31.0 million reads). After adapter trimming and quality filtering, we retained an average of 96.1% of reads (95.4-97%) per library. Subsequently, an average of 24.1 million reads (20.6 to 30 million reads) per library was successfully mapped to the methicillin resistant *Staphylococcus aureus* USA300 reference genome (GenBank accession no. CP000255.1), with an average of only 1.5% of reads (0.9-2.8%) mapping to ribosomal genes. The libraries had an average estimated depth of coverage of 9166-fold (7842x to 11384x), with only 1.7% of genes having <10 mapped reads (2585/2631 genes with sufficient sample size for determining differential expression). For comparisons of TSB versus urine, Euclidean distances between samples (Supplementary Fig S1) and principal-component analysis (Supplementary Fig S2) revealed that expression patterns in biological replicates of each treatment group were more similar to each other than they were to those in biological replicates of the contrasting treatment group. In comparison to the control, MRSA-1369 cultivated in human urine showed significant changes (defined as absolute log_2_ fold change, |log_2_FC| >1 and adjusted P value (P_adj_) of ≤ 0.05) in the expression of 861 genes of which 461 were significantly upregulated (Supplementary Table S1) while 400 were significantly downregulated (Supplementary Table S2). RNASeq results were confirmed by quantitative real-time PCR-based analysis of expression of a panel of 11 overlapping genes. Fold change was strongly correlated across these genes (r^2^ = 0.89; Supplementary Fig S3).

### MRSA-1369 differential gene expression in human urine

Healthy human urine is made of low concentration of amino acids and short peptides, trace amounts of transition metals and more than 2500 metabolites [14, 15]. To examine the metabolic adaptations in MRSA-1369 in response to the nutrient-limiting growth conditions in human urine, we compared the expression of metabolic genes in TSB-control and human urine exposed MRSA-1369. We observed that TCA cycle (GO:0006099) was one of the significantly enriched GO terms (Table 1). More specifically, in comparison to TSB-control, MRSA-1369 in human urine exhibited significantly increased expression of genes encoding enzymes that catalyze reactions in tricarboxylic acid (TCA) cycle such as succinate dehydrogenase (*sdhC, sdhA, sdhB*), succinyl coA synthase (*sucC, sucD*), 2-oxoglutarate dehydrogenase (*sucB, sucA*), isocitrate dehydrogenase (*icd*), aconitate hydratase (*acnA*), citrate synthase (*gltA*), fumarate hydratase (*fumC*), and malate:quinone oxidoreductase (*mqo*). Cultivation in human urine also increased transcription of genes encoding phosphoenolpyruvate carboxykinase (*pckA*) that catalyzes first irreversible step in gluconeogenesis by converting oxaloacetate into phosphoenol pyruvate and pyruvate synthase (*aka* pyruvate ferredoxin oxidoreductase) which catalyzes conversion of pyruvate into acetyl coA, the substrate for TCA cycle (Table 2).

**Table 1:**
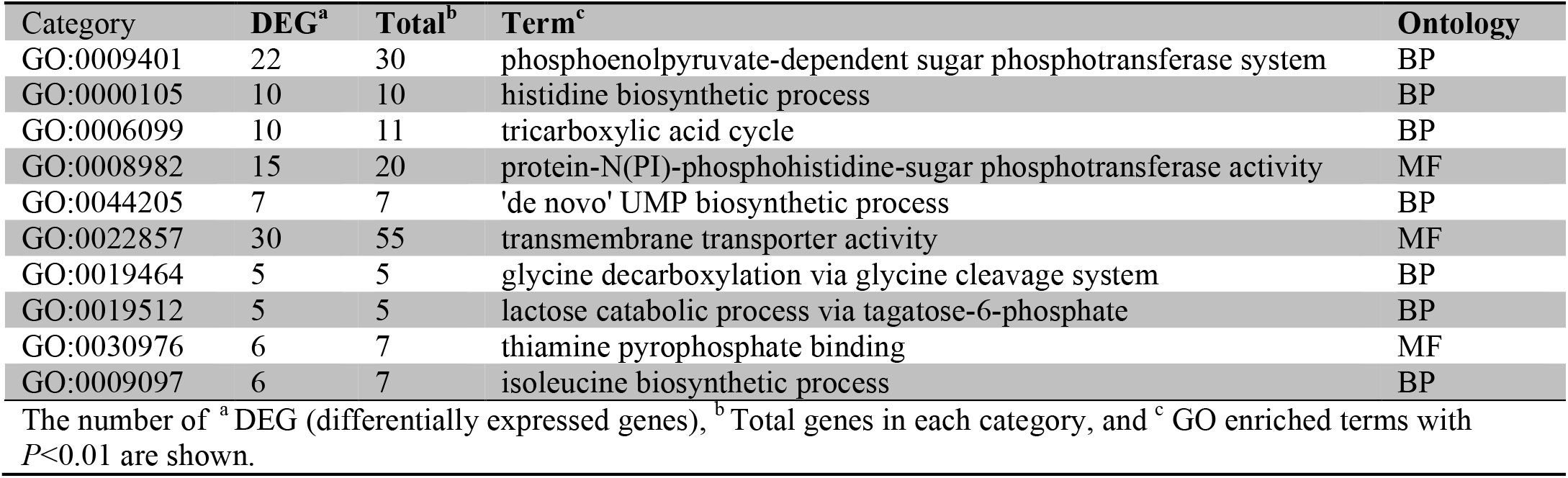
Gene ontology (GO) enrichment analysis

**Table 2:**
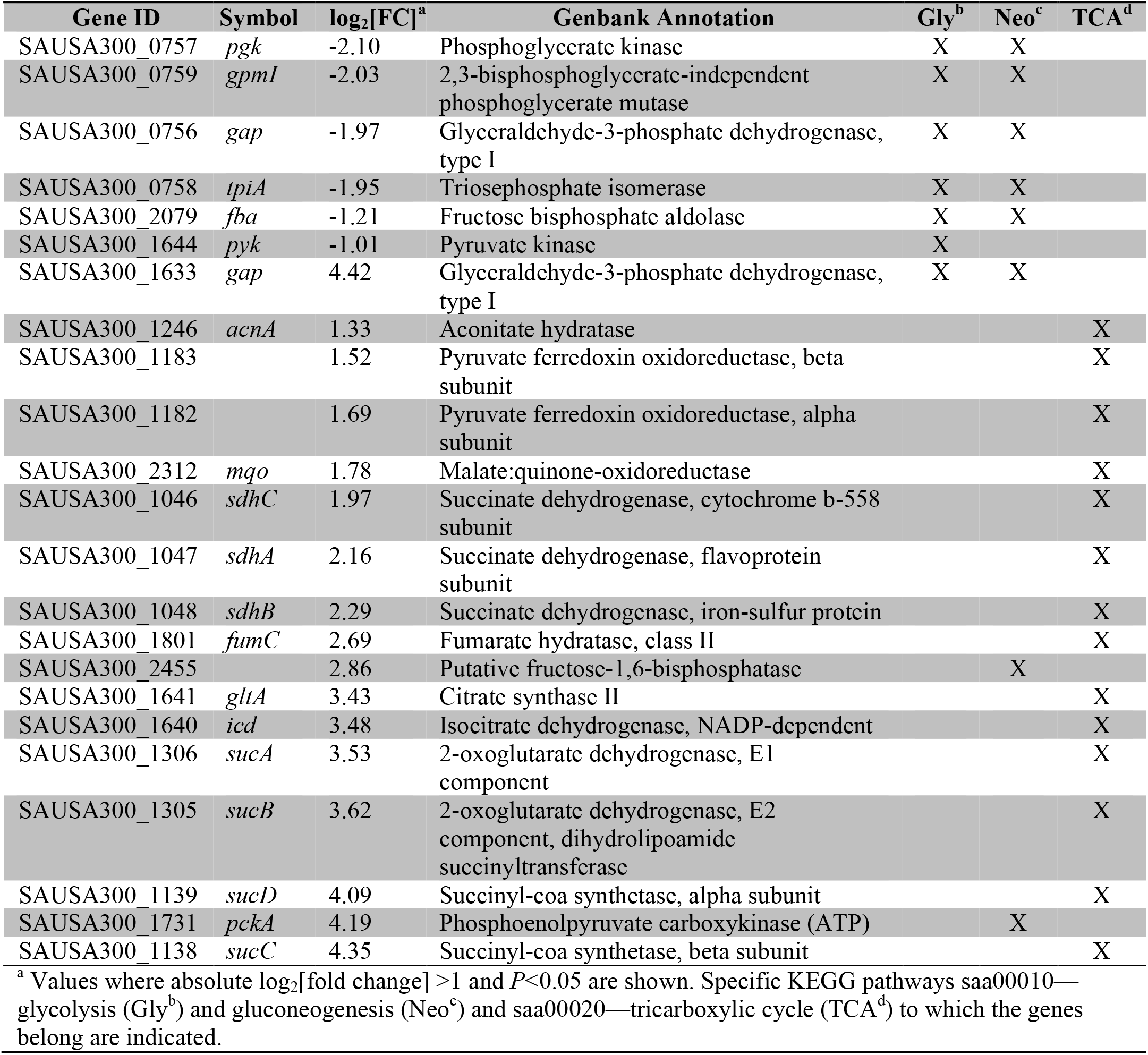
RNA-Seq results for genes encoding enzymes catalyzing glycolysis, gluconeogenesis, and TCA cycle

Given the status of MRSA as a major human pathogen, we were interested in analyzing RNASeq data to define the changes in virulence genes and associated regulators following 2h-long cultivation in human urine. To achieve this objective, we focused on genes categorized into KEGG pathway defined as “*Staphylococcus aureus* infection” (saa05150) and on genes experimentally shown to play a role in MRSA infection (Table 3). In human urine-cultivated MRSA-1369, we observed significant downregulation of genes encoding surface proteins involved in colonization such as *sasG/E* and clumping factor B (*clfB*), cytolytic toxins Panton-Valentine leukocidin (*lukF/S*), superantigen-like proteins *seq* and *eta*. In contrast, human urine significantly upregulated expression of genes encoding surface proteins clumping factor A (*clfA*), fibrinogen-binding protein (*efb*) and cytolytic α-hemolysin precursor (*hla*) and γ-hemolysin (*hlgA*/*B*/*C*). In addition, human urine also significantly altered expression of transcriptional regulators such as *codY, agr* quorum sensing system (*agrDCA*), sarA family regulators (*rot, sarA, mgrA*), two component systems *saeS/R* and *lytR/S*, stress-related gene *ctsR*, and heat-shock proteins *clpPCB, dnaJ/K, grpE*, and *groL/ES* (Table 3).

**Table 3:** RNA-Seq results for virulence genes categorized into KEGG pathway saa05150—“*Staphylococcus aureus* infection” and important transcriptional regulators

| Gene ID | Symbol | log<sub>2</sub>[FC]<sup>a</sup> | Genbank Annotation |
| --- | --- | --- | --- |
| MSCRMM-Colonization |  |  |  |
| SAUSA300_2435 | *sasG_E* | -1.25 | Cell wall surface anchor family protein |
| SAUSA300_2436 | *sasG_E* | -1.51 | Putative cell wall surface anchor family protein |
| SAUSA300_2565 | *clfB* | -1.08 | Clumping factor B |
| Surface Proteins |  |  |  |
| SAUSA300_0772 | *clfA* | 1.27 | Clumping factor A |
| SAUSA300_1055 | *efb* | 3.09 | Fibrinogen-binding protein |
| SAUSA300_1920 | *chs* | -1.33 | Chemotaxis-inhibiting protein CHIPS |
| SAUSA300_1101 |  | 1.24 | Putative fibronectin/fibrinogen binding protein |
| SAUSA300_1052 | *efbC* | 3.19 | Fibrinogen-binding protein |
| Antimicrobial Activity |  |  |  |
| SAUSA300_0647 | *vraF* | -1.07 | CAMP transport system ATP-binding protein |
| SAUSA300_0835 | *dltA* | -1.40 | D-alanine-activating enzyme/D-alanine-D-alanyl |
| Cytolytic Toxins |  |  |  |
| SAUSA300_2365 | *hlgA* | 6.80 | γ-hemolysin component A |
| SAUSA300_2366 | *hlgC* | 3.63 | γ-hemolysin component C |
| SAUSA300_2367 | *hlgB* | 3.14 | γ-hemolysin component B |
| SAUSA300_1058 | *Hla* | 3.52 | α-Hemolysin precursor |
| Superantigens/ Superantigen like proteins |  |  |  |
| SAUSA300_0801 | *Seq* | -1.04 | Staphylococcal enterotoxin Q |
| SAUSA300_1065 | *Eta* | -1.52 | Exfoliative toxin A |
| **Transcriptional Regulators and two component systems** |  |  |  |
| SAUSA300_0254 | *lytS* | -1.59 | Sensor histidine kinase |
| SAUSA300_0255 | *lytR* | -2.29 | Sensory transduction protein |
| SAUSA300_0507 | *ctsR* | 3.15 | Transcriptional regulator |
| SAUSA300_0605 | *sarA* | 1.51 | Staphylococcal accessory regulator A |
| SAUSA300_0672 | *mgrA* | -1.95 | Transcriptional regulator, marr family |
| SAUSA300_0690 | *saeS* | 2.13 | Sensor histidine kinase, |
| SAUSA300_0691 | *saeR* | 2.49 | DNA-binding response regulator |
| SAUSA300_1148 | *codY* | 1.45 | GTP-sensing transcriptional pleiotropic repressor |
| SAUSA300_1514 | *fur* | 1.26 | Ferric uptake regulation protein |
| SAUSA300_1542 | *hrcA* | 3.58 | Heat-inducible transcription repressor |
| SAUSA300_1708 | *rot* | 1.98 | Staphylococcal accessory regulator |
| SAUSA300_1990 | *agrD* | 1.09 | Accessory gene regulator protein D |
| SAUSA300_1991 | *agrC* | 1.02 | Accessory gene regulator protein C |
| SAUSA300_1992 | *agrA* | 1.05 | Accessory gene regulator protein A |

<sup>a</sup> Only values where absolute log<sub>2</sub>[fold change] >1 and *P*<0.05 are shown.
Positive values indicate greater expression in the human urine treatment, and negative values indicate greater expression in the TSB-control.

Human urine also induced significantly higher expression of MRSA-1369 genes in pathways for acquisition and export of metals such as iron, nickel, and zinc, which are essential nutrients. The expression of zinc transporters *znuB/C*, SAUSA300_2315, cobalt-nickel transporters *cntFDCB, opp1A*, and *oppBCDFAA* was upregulated in MRSA-1369 exposed to human urine for 2h (Table 4). Human urine also induced upregulation of *sbnABCDEFGHI* and *sirA/B* involved in the production and import of siderophore staphylophorin B, respectively, *fhuC/B* involved in siderophore transport, *isdCDEF*, srtB from iron surface determinant *isd* heme acquisition system, *sstABCD* from catechol/catecholamine iron transporter system, and *htsCBA* importer of staphylosporin A (Table 5, [16, 17]).

**Table 4:** RNA-Seq results for genes categorized into KEGG pathway saa02010—“ABC transporters”

| Gene ID | Symbol | log<sub>2</sub>[FC]<sup>a</sup> | Genbank Annotation |
| --- | --- | --- | --- |
| Mineral and Organic Iron transporter |  |  |  |
| SAUSA300_0219 |  | -1.345 | Putative iron compound A C transporter, iron compound-binding protein |
| SAUSA300_0999 | *potA* | 2.17 | Spermidine/putrescine ABC transporter, ATP-binding protein |
| SAUSA300_1000 | *potB* | 2.48 | Spermidine/putrescine ABC transporter, permease protein |
| SAUSA300_1001 | *potC* | 1.92 | Spermidine/putrescine ABC transporter, permease protein |
| SAUSA300_1002 | *potD* | 1.05 | Spermidine/putrescine ABC transporter, spermidine/putrescine-binding protein |
| Oligosaccharide, polyol, and lipid transport |  |  |  |
| SAUSA300_0209 |  | 1.78 | Putative maltose ABC transporter, maltose-binding protein |
| SAUSA300_0210 |  | 1.89 | Maltose ABC transporter, permease protein |
| SAUSA300_0211 |  | 1.73 | Maltose ABC transporter, permease protein |
| SAUSA300_0263 | *rbsD* | 1.99 | Ribose permease |
| Phosphate and amino acid transport |  |  |  |
| SAUSA300_1280 | *pstB* | 1.21 | Phosphate ABC transporter, ATP-binding protein |
| Cysteine transporter |  |  |  |
| SAUSA300_2357 |  | -1.24 | ABC transporter, ATP-binding protein |
| SAUSA300_2358 |  | -1.11 | ABC transporter, permease protein |
| SAUSA300_2359 |  | -1.19 | Amino acid ABC transporter, amino acid-binding protein |
| Oligopeptide transport |  |  |  |
| SAUSA300_0887 | *oppB* | 1.11 | Oligopeptide ABC transporter, permease protein |
| SAUSA300_0888 | *oppC* | 1.25 | Oligopeptide ABC transporter, permease protein |
| SAUSA300_0889 | *oppD* | 1.36 | Oligopeptide ABC transporter, ATP-binding protein |
| SAUSA300_0890 | *oppF* | 1.42 | Oligopeptide ABC transporter, ATP-binding protein |
| SAUSA300_0891 | *oppA* | 1.41 | Oligopeptide ABC transporter, substrate-binding protein |
| SAUSA300_0892 | *oppA* | 1.20 | Oligopeptide ABC transporter, oligopeptide-binding protein |
| Nickel transport |  |  |  |
| SAUSA300_2407 | *cntF* | 1.70 | Oligopeptide ABC transporter, ATP-binding protein |
| SAUSA300_2408 | *cntD* | 1.90 | Oligopeptide ABC transporter, ATP-binding protein |
| SAUSA300_2409 | *cntC* | 1.89 | Oligopeptide ABC transporter, permease protein |
| SAUSA300_2410 | *cntB* | 2.80 | Oligopeptide ABC transporter, permease protein |
| SAUSA300_2411 | *opp-1A* | 2.99 | Oligopeptide permease, peptide-binding protein |
| Iron Siderophore transporter |  |  |  |
| SAUSA300_2134 |  | 1.49 | Iron compound ABC transporter, permease protein |
| SAUSA300_2135 |  | 1.70 | Iron compound ABC transporter, permease protein |
| SAUSA300_2136 |  | 2.86 | Iron compound ABC transporter, iron compound-binding protein |
| Zinc transporter |  |  |  |
| SAUSA300_1515 |  | 1.47 | ABC transporter, permease protein |
| SAUSA300_1516 |  | 1.76 | ABC transporter, ATP-binding protein |
| SAUSA300_2351 |  | 1.94 | Zn-binding lipoprotein adca-like protein |
| Biotin transporter |  |  |  |
| SAUSA300_0977 |  | -2.49 | Cobalt transport family protein |
| Unclassified |  |  |  |
| SAUSA300_1760 | *epiG* | 2.18 | Lantibiotic epidermin immunity protein F |
| SAUSA300_1761 | *epiE* | 1.76 | Lantibiotic epidermin immunity protein F |
| SAUSA300_1762 | *epiF* | 1.39 | Lantibiotic epidermin immunity protein F |
| SAUSA300_2465 |  | -1.41 | ABC transporter, ATP-binding protein |
| SAUSA300_0647 |  | -1.07 | ABC transporter, ATP-binding protein |
| SAUSA300_0672 |  | -1.95 | Transcriptional regulator, marR family |

<sup>a</sup> Only values where absolute log<sub>2</sub>[fold change] >1 and *P* < 0.05 are shown.
Positive values indicate greater expression in the human urine treatment, and negative values indicate greater expression in the TSB-control.

**Table 5:** RNA-Seq results for genes involved in iron transport

| Gene ID | Symbol | log<sub>2</sub>[FC]<sup>a</sup> | Genbank Annotation |
| --- | --- | --- | --- |
| SAUSA300_0633 | *fhuA* | 1.83 | Ferrichrome transport ATP-binding protein fhua |
| SAUSA300_0634 | *fhuB* | 1.08 | Ferrichrome transport permease protein fhub |
| SAUSA300_0116 | *sirB* | 1.84 | Iron compound ABC transporter, permease protein sirb |
| SAUSA300_0117 | *sirA* | 2.30 | Iron compound ABC transporter, iron compound-binding protein sira |
| SAUSA300_0118 | *sbnA* | 2.64 | Pyridoxal-phosphate dependent enzyme superfamily |
| SAUSA300_0119 | *sbnB* | 2.29 | Ornithine cyclodeaminase |
| SAUSA300_0120 | *sbnC* | 2.16 | Siderophore biosynthesis protein, iucc family |
| SAUSA300_0121 | *sbnD* | 1.90 | Putative drug transporter |
| SAUSA300_0122 | *sbnE* | 1.54 | Siderophore biosynthesis protein, iuca/iucc family |
| SAUSA300_0123 | *sbnF* | 1.37 | Siderophore biosynthesis protein, iucc family |
| SAUSA300_0124 | *sbnG* | 1.42 | HPCH/HPAI aldolase family protein |
| SAUSA300_0125 | *sbnH* | 1.27 | Pyridoxal-dependent decarboxylase |
| SAUSA300_0126 | *sbnI* | 1.03 | Conserved hypothetical protein |
| SAUSA300_0718 | *sstA* | 2.45 | Iron compound ABC transporter, permease |
| SAUSA300_0719 | *sstB* | 2.52 | Iron compound ABC transporter, permease protein |
| SAUSA300_0720 | *sstC* | 2.88 | Putative iron compound ABC transporter, ATP-binding protein |
| SAUSA300_0721 | *sstD* | 3.77 | Transferrin receptor |
| SAUSA300_1030 | *isdC* | 2.59 | Iron transport associated domain protein |
| SAUSA300_1031 | *isdD* | 2.01 | Conserved hypothetical protein |
| SAUSA300_1032 | *isdE* | 2.31 | Putative iron compound ABC transporter, iron compound-binding protein |
| SAUSA300_1033 | *isdF* | 1.39 | Iron/heme permease |
| SAUSA300_1034 | *srtB* | 1.64 | Sortase B |
| SAUSA300_1035 | *isdG* | 1.11 | Conserved hypothetical protein |
| SAUSA300_2134 | *htsC* | 1.49 | Iron compound ABC transporter, permease protein |
| SAUSA300_2135 | *htsB* | 1.70 | Iron compound ABC transporter, permease protein |
| SAUSA300_2136 | *htsA* | 2.86 | Iron compound ABC transporter, iron compound-binding protein |

<sup>a</sup> Only values where absolute log<sub>2</sub>[fold change] >1 and *P*<0.05 are shown.
Positive values indicate greater expression in the human urine treatment, and negative values indicate greater expression in the TSB-control.

## Discussion

An overarching objective of our research is to delineate the effects of growth limiting and hostile microenvironment of the urinary tract on the physiology of various bacterial pathogens. In line with this objective, for this report we examined virulence characteristics and gene expression of various *S. aureus* strains using human urine as a culture medium to mimic conditions encountered during the colonization of human urinary tract. The specific strains analyzed in this study include two urinary isolates, MRSA-1369 and PUTS-1 and a prototypical MRSA strain USA300, all of which were able to grow in human urine. Although PUTS-1, a clinical isolate from a woman with asymptomatic bacteriuria exhibited higher doubling time compared to the MRSA strains. This matches previous work showing that asymptomatic bacteriuria strains of uropathogenic *E. coli* and *Streptococcus agalactiae* exhibit rapid growth compared to those causing cystitis [18, 19]. Our observations further suggest that the induction of specific virulence characteristics by human urine is strain-dependent. In addition, we also compared the global transcriptome profiles of uropathogenic strain MRSA-1369 exposed *in vitro* to either human urine or to nutrient-rich culture medium TSB (control). The RNASeq observations constitute the first step in defining molecular basis for results from virulence assays as discussed below.

*S. aureus* strains infecting humans secrete numerous cytolytic exotoxins such as α-hemolysin and leukotoxins as well as cytolytic peptides called phenol soluble modulins, which play an important role in staphylococcal pathogenesis by facilitating tissue damage, immune evasion, and dissemination [12]. We observed upregulation of *hla* encoding α-hemolysin and *hlgACB* cluster encoding γ-hemolysin, both of which are known to target a broad range of host cells including red blood cells [20, 21]. The *hla* gene is expressed by most clinical isolates of MRSA and its expression level is correlated with the disease severity [22]. Moreover, α-hemolysin is known to be essential for MRSA virulence in animal models of skin and soft tissue infections (SSTI), pneumonia, and bacteremia [20]. The γ-hemolysin belongs to the leukotoxin family of cytolysins, which are made of two different protein components that assemble to form a β barrel pores in the host plasma membrane. The γ-hemolysin is highly prevalent in *S. aureus* clinical isolates from human nose and blood [21]. Although, it must be noted that, while *hlgACB* genes are upregulated in human blood, the contribution of γ-hemolysin to staphylococcal virulence is modest as shown by the results from infection experiments comparing WT MRSA with Δ*hlgACB* mutant defective in γ-hemolysin production in animal models of eye infection[23], bacteremia[24], and septic arthritis [25]. The increased expression of *hla* and *hlgACB* in human urine and corresponding increase in hemolytic activity suggests that either one or both toxins play an important role in the urinary pathogenesis of MRSA-1369. Moreover, the absence of hemolytic activity in PUTS-1 may be linked to the reduced level of *hla* expression in human urine. It will be worthwhile for the future research to investigate whether one or more genes encoding specific cytolytic toxins is inactivated in PUTS-1 and whether its comparatively rapid growth would compensate for the lack of hemolysis and offer it a competitive advantage in asymptomatic colonization of the urinary tract.

Multiple nonspecific and specific mechanisms of adherence facilitate early steps of staphylococcal colonization of host tissue by promoting bacterial attachment to host cells and extracellular matrix proteins. Non-specific mechanisms include changes in surface charge and hydrophobicity, while specific mechanisms include number of cell wall anchored adhesins (MSCRAMMs—Microbial Surface Components Recognizing Adhesive Matrix Molecules), which specifically adhere to ECM proteins collagen, fibrinogen, and fibronectin [26]. Adherence to bladder epithelium is the first important step in urinary pathogenesis as it prevents removal of uropathogens by the gush of urine. Human urine specifically induced adherence to human bladder epithelial cells by MRSA-1369 and USA300. The hydrophobicity was significantly higher in MRSA-1369 cultivated in human urine. Based on gene expression data, the increased adherence in MRSA strains cultivated in human urine could potentially be mediated by the product of one or more of the surface adhesin genes *clfA, efb, efb-c*, and SAUSA300_1101. In contrast, binding to fibronectin was reduced in all three staphylococcal strains. This matches with gene expression results (qRTPCR and/or RNASeq) showing that that the expression of genes encoding principal fibronectin adhesins, fibronectin binding protein (*fnbAB*) was downregulated in human urine in MRSA-1369, PUTS-1, and USA300.

Clumping factor A and B (clfA, clfB) are important mediators of MRSA binding to fibrinogen [27]. MRSA adherence to fibrinogen is implicated in the pathogenesis of CAUTI as MRSA infection is shown to increase fibrinogen deposition on catheters, and because Δ*clfB* deletion mutant in MRSA-1369 shows reduced bacterial burden on catheter implant in a mouse model of CAUTI [8]. Interestingly, deletion of *clfA* (Δ*clfA*) does not affect bacterial burden on catheter implant compared to WT MRSA1369 [8]. In the light of this information, how our observations that human urine mediates upregulation of *clfA* and downregulation of *clfB* expression may shape MRSA uropathogenesis warrant further experimentation in a mouse model.

Primarily made of urea, inorganic salts, creatinine, organic acids, small quantities of amino acids, trace amounts of transition metals and other water-soluble waste products from blood, the urine is a nutrient-poor culture medium. To colonize urinary tract, uropathogens must catabolize amino acids via tricarboxylic acid (TCA) cycle to provide substrates for gluconeogenesis. The increased expression of genes encoding TCA cycle/gluconeogenesis enzymes and oligopeptide transporters (*oppBCDFAA*) when MRSA-1369 was cultivated in human urine implicates these central metabolic pathways in MRSA uropathogenesis. This is similar to previous reports showing that UPEC mutants ablated in oligopeptide and dipeptide transport as well as mutants lacking enzymes catalyzing TCA cycle or gluconeogenesis exhibit fitness defects in the mouse model of ascending UTI [28]. In addition, our results also implicate transporter systems for nickel, iron and zinc in MRSA uropathogenesis. Both bacterial pathogens and their eukaryotic hosts require transition metals for survival; hosts exert a tight control over metal homeostasis as a defense mechanism against infections while bacteria produce acquisition (iron siderophores) and export systems to chelate metals from the host. Human urine contains trace quantities of iron (0.089µM/mM of creatinine), nickel (0.0080µM/mM of creatinine), cobalt (0.0014 µM/mM of creatinine), and zinc (0.46µM/mM of creatinine) [15].

Most of the research so far is focused on revealing the interactions between host immunity and MRSA virulence in the context of predominant MRSA infections of skin, soft-tissue and lungs. In contrast, MRSA-host interactions within the urinary environment are largely unexplored. To survive in the urinary niches, MRSA must rapidly adapt to unique challenges in the form of nutrient unavailability, mobilization of immune defenses, acidic pH, osmolarity, and shear stress due to urine flow. In this report, we have correlated alterations in virulence characteristics visualized by *in vitro* assays with changes in the expression of specific genes, which, we are aware, is only the first step in understanding MRSA physiology in the urinary tract. In the future, such a correlation should be confirmed by comparing deletion mutants targeting specific genes with WT MRSA by *in vitro* virulence assays and in a mouse model of ascending UTI. We will be remiss if we do not mention important caveats of this study that changes in mRNA transcripts levels do not guarantee corresponding changes in protein levels and that our results provide but a snapshot into the transcriptome and virulence of MRSA at 2h time point following cultivation in human urine. To address these caveats, the future research work should focus on the analysis of host/MRSA transcriptomes and proteomes in infected murine bladder and kidney tissues harvested at different time points.

## Materials and methods

### Bacterial strains and culture conditions

*Staphylococcus aureus* strains used in this study are MRSA-1369, PUTS-1, and USA300 (Table 6, [8, 29]). On the day of experiment, overnight cultures in TSB at 37°C and shaking at 200 rpm were diluted 1:10 and grown to OD_600_ of 0.6. These are referred as mid-log cultures.

**Table 6:** Strains used in this study

| Strain | Isolation site | Resistance | Reference cited |
| --- | --- | --- | --- |
| MRSA 1369 | Urine | Methicillin | [8] |
| PUTS-1 4-147 | Urine, Asymptomatic | Penicillin | [8] |
| USA300 | Skin and soft tissue | Methicillin | [29] |

Urine from healthy female volunteers (protocol approved by UL Lafayette IRB) was collected after informed consent was obtained from each volunteer. Urine was immediately filtered-sterilized using a 0.22µm filter and stored in 1.5ml aliquots at −80°C. At the time of experiments, urine aliquots from three-five different donors were warmed to 37°C and mixed. Mid-log bacterial cultures were exposed to either TSB (control) or human urine for 2 h at 37°C. Bacteria were then centrifuged, washed in Dulbecco’s phosphate-buffered saline (D-PBS) and used in various *in vitro* virulence assays and for RNA extractions as specified below. To enumerate the colony forming units (CFU)/ml we plated serial, 10-fold dilutions of bacteria on TS Agar or CHROMagar™.

### Growth curve

*S. aureus* strains were cultivated in TSB or human urine at 37°C without shaking. CFUs were enumerated by dilution plating at different time points up to 24h.

### *In vitro* adherence assay

Approximately 2 ×10^5^ cells of 5637 human bladder epithelial cell line (ATCC HTB-9) maintained in RPMI supplemented with 10% fetal bovine serum (FBS) were seeded per well in 6 well cell culture plate and incubated at 37°C in 5% CO_2_ to get confluent monolayers. The monolayers were weaned (grown in plain medium without FBS) for 24 h before infection. The monolayers were infected at an MOI (multiplicity of infection) of 10 with MRSA strains either in the presence of plain RPMI (control) or human urine. Plates were centrifuged to facilitate contact between MRSA and bladder cells. After incubation at 37°C for 2h and 5% CO_2_, supernatant was collected in sterile microcentrifuge tube to determine non-adherent CFUs by dilution plating. Monolayers were then washed 3 times with sterile PBS (containing Ca^++^/ Mg^++^) to remove non-adherent bacteria. Adherent bacteria collected in sterile PBS containing 0.1% Triton-X 100 were enumerated by dilution plating [30].

%Adherence= [Adherent CFU ÷ (Adherent CFU + Supernatant CFU)] x 100

### Fibronectin binding assay

Assay to determine binding of MRSA to human fibronectin was performed as described previously [31]. In brief, a 96-well microtiter plate was coated overnight with 2-fold dilutions (20, 10, 5, 2.5 and 1.25µg/ml) of 100µl of human fibronectin at 4°C. The wells were washed three-times with 0.05% Tween 20 in sterile PBS and then blocked with 100µl of 1% bovine serum albumin solution for 1 h at 37°C. The wells were washed again and 100µl of MRSA (pre-exposed to human urine or TSB control, adjusted to OD_600_=0.45) was added to duplicate wells. After incubation for 2 h at 37°C, non-adherent bacteria were removed by washing. The adherent bacteria were fixed with 100µl of 25% formaldehyde for 30 min and stained with 100µl of 1% crystal violet for 15 min at room temperature. After washing with water and drying (37°C, 2h) crystal violet was extracted with 70%-10% ethanol-methanol mixture. The absorbance was measured at 590 nm.

### Hydrophobicity test

Hydrophobicity was determined using MATH (microbial adhesion to hydrocarbon) assay, wherein *S. aureus* strains pre-exposed to human urine or TSB were centrifuged, resuspended in sterile DPBS, and adjusted to OD_600_=0.6. One ml of each bacterial suspension was mixed with 125 µl hexadecane by vortexing for 1 min and incubated at room temperature for 30 min [32]. The CFU/ml before addition of hexadecane (C_i_) and CFU/ml in the aqueous phase (C_aq_) after incubation with hexadecane were enumerated by dilution plating.

% Hydrophobicity=[(C_i_-C_aq_) ÷ C_i_] x100

### Antimicrobial peptide (AMP) resistance Assay

To determine the sensitivity of bacteria to human AMP, cathelicidin LL-37, *S. aureus* strains pre-exposed to TSB or human urine were washed, resuspended in sterile DPBS supplemented with 20% TSB either without or with 50 µM LL-37 for 1h. Bacteria were enumerated by dilution plating. % AMP resistance = (CFU_treated_/ CFU_untreated_) x 100.

### Hemolysis assay

Staphylococcal strains pre-exposed to TSB (control) or human urine used to generate two-fold serial dilutions starting from 10^8^CFU/ml. Dilutions were mixed with equal volume of 1% sheep erythrocytes in PBS in 96-well conical bottom plate and incubated at 37°C. After 2 h, unlysed RBCs were pelleted at 3000 rpm for 10 minutes and 100µl supernatant transferred to a fresh plate. Absorbance at 420nm was measured to estimate hemoglobin release [33]. A_420_ readings for RBCs treated with PBS or 0.1% Triton-X 100 were used to define baseline (A_NC_) and 100% hemolysis (A_PC_), respectively. % Hemolysis = [(A_treatment_-A_NC_) ÷ (A_PC_ - A_NC_)] x 100.

### Autolysis assay

Staphylococci pre-exposed to TSB or HU were resuspended in 0.2% triton X-100 in sterile PBS and incubated at 37°C without shaking [34]. We measured initial OD_600_ (OD_i_) at the beginning of the experiment and then at 1h intervals for 4 hours (OD_t_).

% Autolysis = [(OD_i_ – OD_t_)/ OD_i_] x 100

### RNA extraction and quantitative real-time PCR (qRT-PCR)

*S. aureus* strains in the exponential phase of growth were exposed in TSB, human urine for 2h at 37°C. Bacterial RNA was extracted using Ambion® Ribopure™ kit (Thermofisher) and quantified using Synergy™ HTX Multi-Mode Microplate Reader (Biotek). Next, 1µg RNA was reverse transcribed using the high-capacity cDNA reverse transcription kit (Applied Biosystems™). Quantitative real-time PCR (qRT-PCR) was carried out using SYBR green master mix (Applied Biosystems™) in a StepOne Plus thermal cycler (Applied Biosystems). RQ values were calculated by comparing ΔΔC_T_ value. List of the primers used for qRT-PCR is shown in table 7.

**Table 7:**
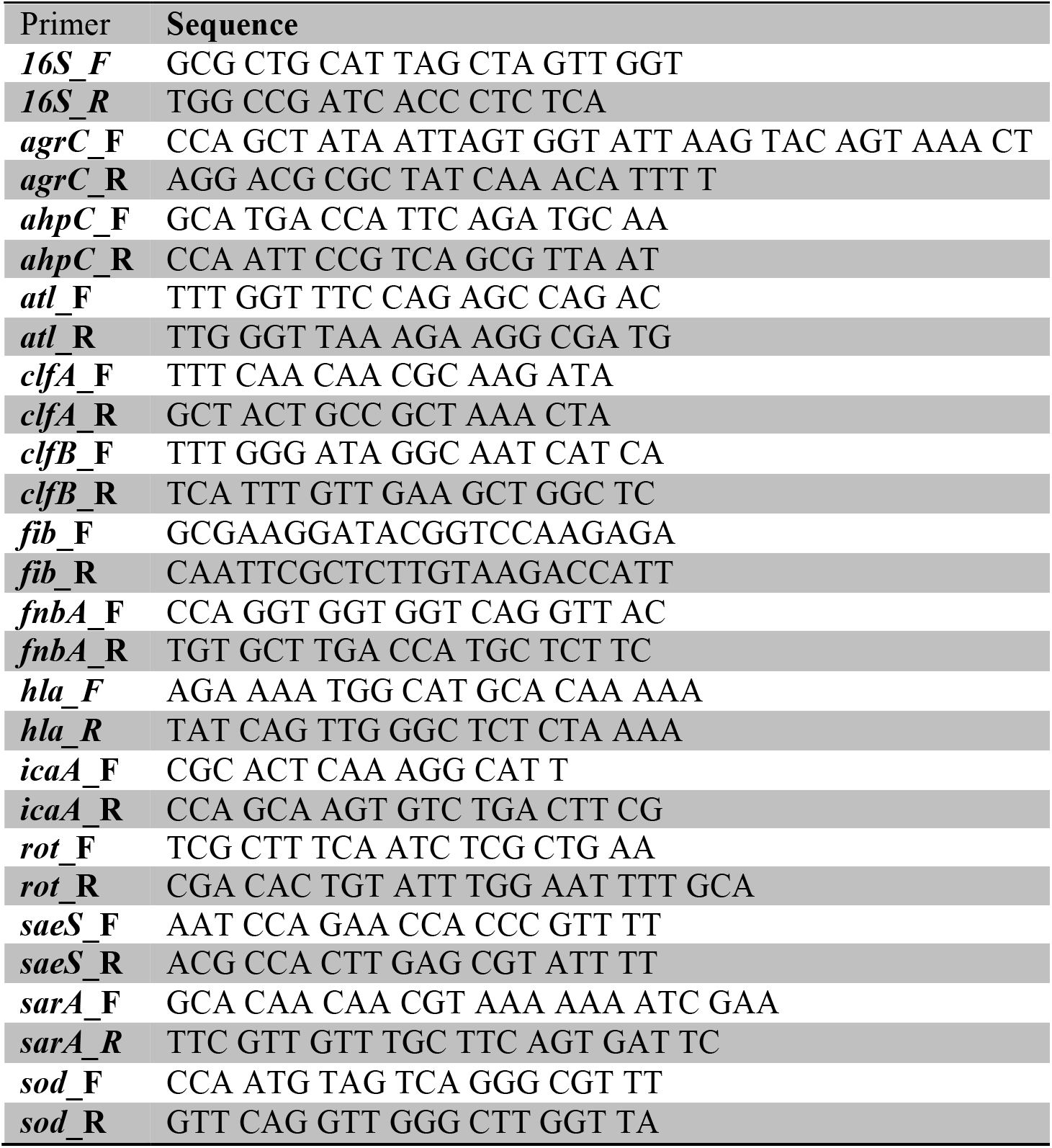
Sequences of primers used for qRTPCR in this study

### Statistical analysis

Data from multiple replicates for each experiment are pooled together. Graphing and statistical analyses are done using GraphPad Prism 9 software. Results are expressed as the means ± standard deviation from data collected from two or more biological replicates. In the case of counts data (CFU/ml), not normally distributed, Mann-Whitney U statistic is used to evaluate the difference between two groups. The difference between groups is considered significant if *P* ≤ 0.05.

### RNASeq data analysis

RNA for RNASeq was extracted as above. All library construction and initial analysis of differential expression was done by GENEWIZ (New Jersey, USA). Library construction included DNase treatment (TURBO™ DNase, ThermoFisher Scientific) and rDNA depletion (QIAseq FastSelect, Qiagen) followed by RNA fragmentation and random priming. cDNA synthesis (NEBNext® Ultra™ II, New England Biolabs) was followed by end repair, 5’ phosphorylation and dA-tailing before. Libraries were sequenced on a partial lane of Illumina HiSeq 400 with 150bp PE sequencing. Quality of sequence data was assessed using FastQC. All reads were quality filtered and trimmed using Trimmomatic v 0.36 with default settings [35]. Reads were mapped to *Staphylococcus aureus subsp aureus* USA300_FPR3757 genome using bowtie v 2.2.6 [36]and hit count for individual genes were generated using the featurecounts common in the subreads package v 1.5.2 [37]. Genes with <10 reads were dropped from the analysis for differential expression. Differential expression for each gene was assessed using Wald tests implemented in DESeq2 [38]. Genes with an adjusted p-value < 0.05 and absolute log_2_ fold change > 1 were categorized as differentially expressed genes. GO enrichment analyses were conducted using the goseq v package in R [39]. Gene ontology (GO) terms and gene lengths were extracted for each gene in the *Staphylococcus aureus subsp. aureus* USA300_FPR3757 from the UniProt website [40]. We determined whether DEGs between treatments were significantly overrepresented within molecular function, biological process, and cellular component GO terms using a Wallenius approximation and accounting for gene length bias using a probability weight function. Because of the inherent difficulties with multiple testing and correcting for multiple testing in GO analyses, we simply consider any term with p < 0.01 as statistically significant. KEGG pathway enrichment analyses were conducted using the KEGGREST Bioconductor package v [41] and a custom script. KEGGREST was used to download lists of pathways and genes within pathways from the KEGG website for *Staphylococcus aureus subsp. aureus* USA300_FPR3757 (organism code *saa*). We assessed whether DEGs are overrepresented in certain pathways by using a Wilcoxon rank-sum test to determine whether p-values for genes within a focal pathway are less than the p-values of genes that are not within the pathway.

---

**Supplementary Figure S1:**
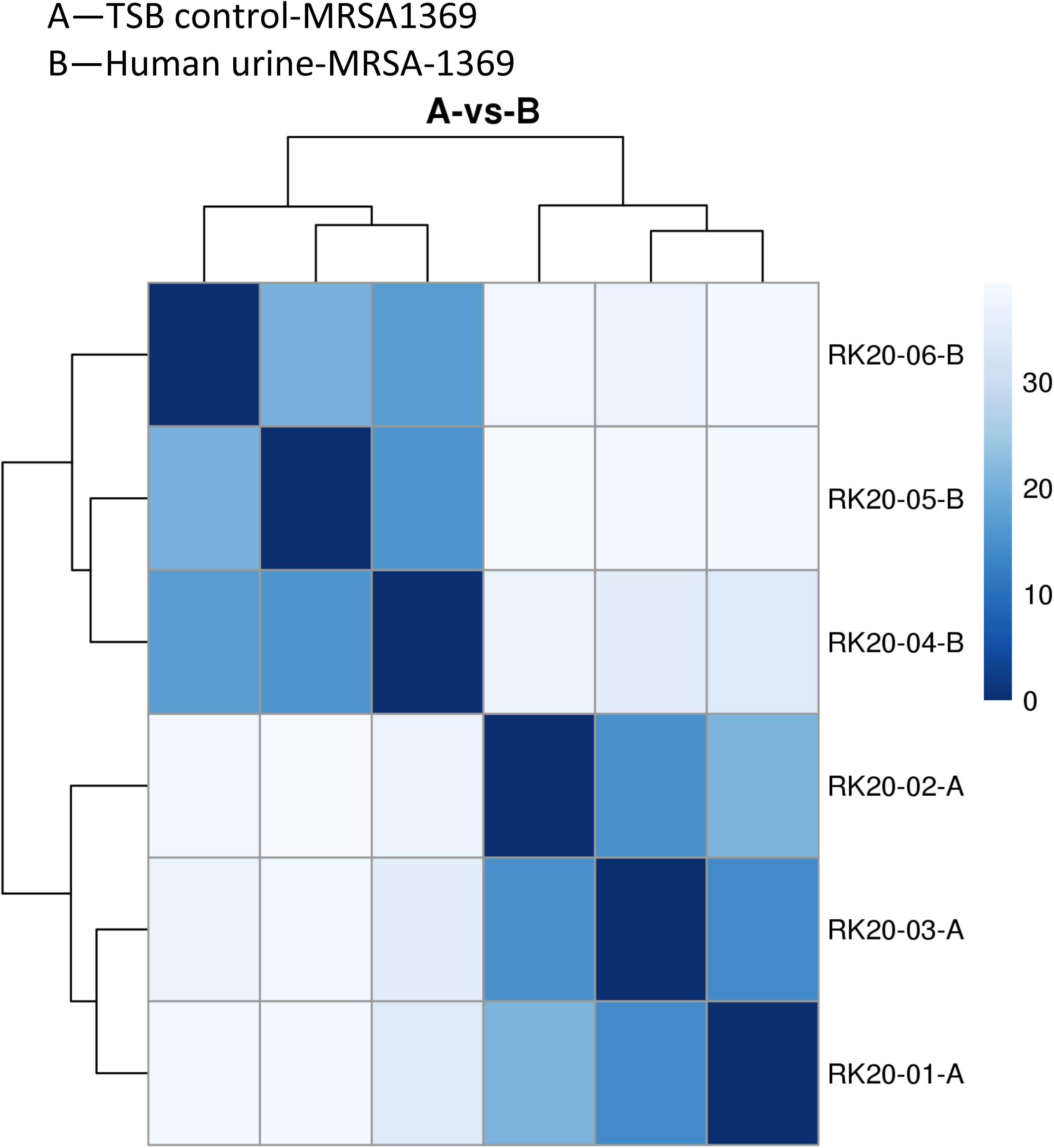
Distances measured using expression values from TSB-control (A) and human urine-exposed (B) MRSA-1369.

**Supplementary Figure S2:**
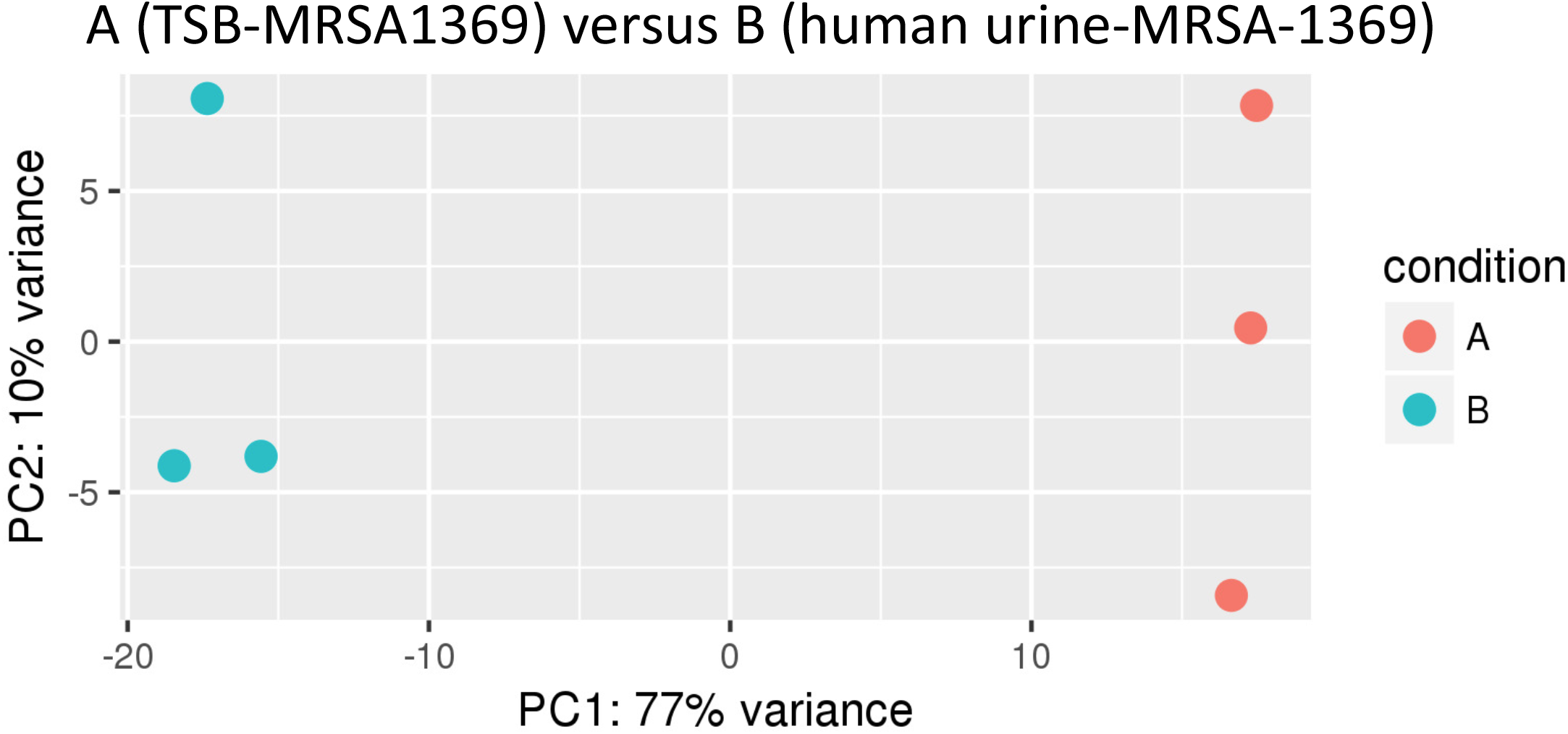
Principal components analysis to reveal similarity within and between groups A (TSB-control) and B (human urine-exposed MRSA-1369).

**Supplementary Figure S3:**
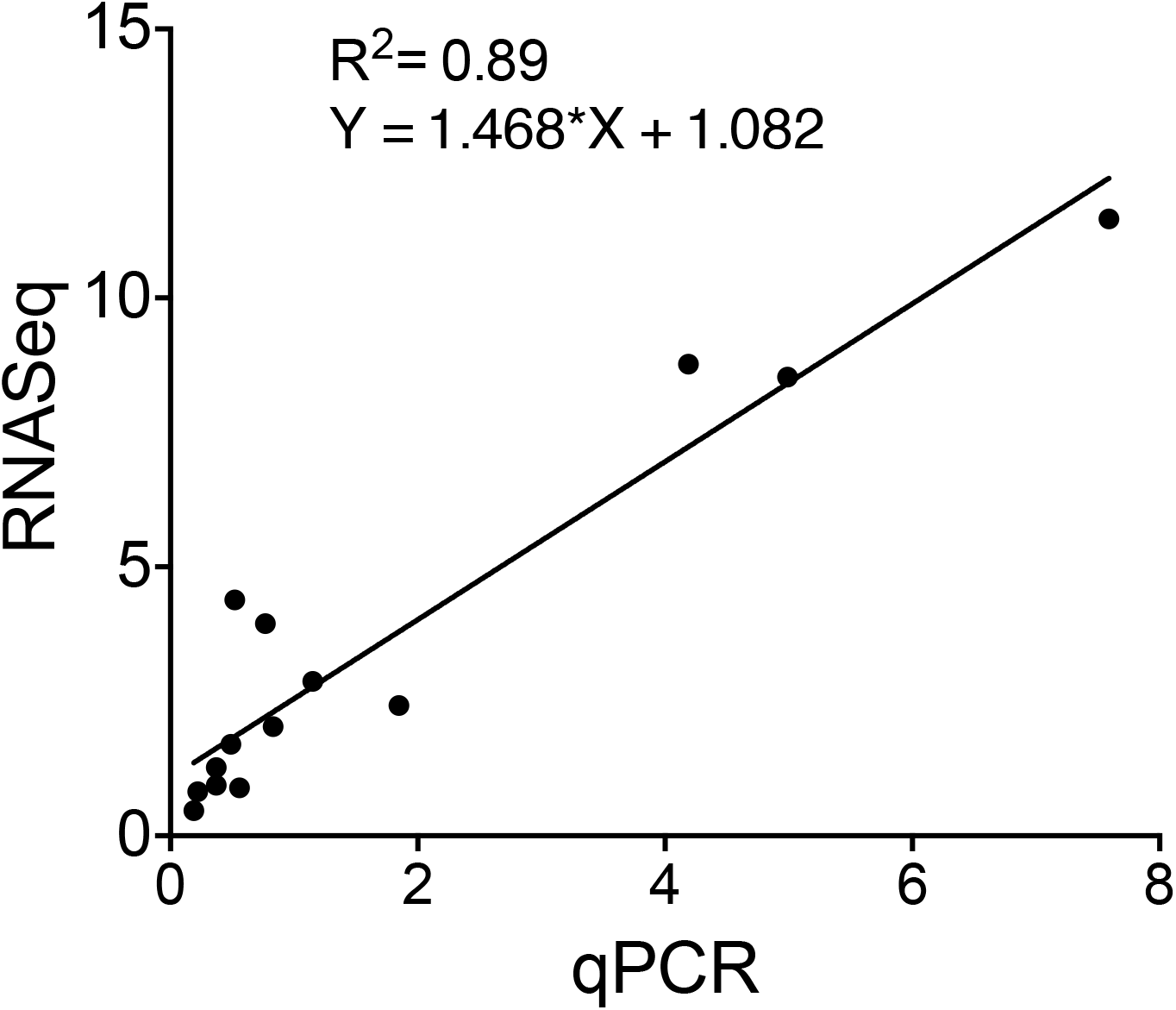
Correlation between RNASeq and qRTPCR results.

## References

1. Medina, M. and E. Castillo-Pino, An introduction to the epidemiology and burden of urinary tract infections. Ther Adv Urol, 2019. 11: p. 1756287219832172.

2. Bishara, J., et al., Healthcare-associated vs. hospital-acquired Staphylococcus aureus bacteremia. Int J Infect Dis, 2012. 16(6): p. e457–63.

3. Stokes, W., et al., Incidence and Outcomes of Staphylococcus aureus Bacteriuria: A Population-based Study. Clin Infect Dis, 2019. 69(6): p. 963–969.

4. Karakonstantis, S. and D. Kalemaki, Evaluation and management of Staphylococcus aureus bacteriuria: an updated review. Infection, 2018. 46(3): p. 293–301.

5. Gad, G.F., et al., Detection of icaA, icaD genes and biofilm production by Staphylococcus aureus and Staphylococcus epidermidis isolated from urinary tract catheterized patients. J Infect Dev Ctries, 2009. 3(5): p. 342–51.

6. Remy, L., et al., The Staphylococcus aureus Opp1 ABC transporter imports nickel and cobalt in zinc-depleted conditions and contributes to virulence. Mol Microbiol, 2013. 87(4): p. 730–43.

7. Hiron, A., et al, A nickel ABC-transporter of Staphylococcus aureus is involved in urinary tract infection. Mol Microbiol, 2010. 77(5): p. 1246–60.

8. Walker, J.N., et al., Catheterization alters bladder ecology to potentiate Staphylococcus aureus infection of the urinary tract. Proc Natl Acad Sci U S A, 2017. 114(41): p. E8721–E8730.

9. Carrel, M., E.N. Perencevich, and M.Z. David, USA300 Methicillin-Resistant Staphylococcus aureus, United States, 2000-2013. Emerg Infect Dis, 2015. 21(11): p. 1973–80.

10. Rosenberg, M., D. Gutnick, and E. Rosenberg, Adherence of Bacteria to Hydrocarbons -a Simple Method for Measuring Cell-Surface Hydrophobicity. Fems Microbiology Letters, 1980. 9(1): p. 29–33.

11. Clarke, S.R., et al., The Staphylococcus aureus surface protein IsdA mediates resistance to innate defenses of human skin. Cell Host Microbe, 2007. 1(3): p. 199–212.

12. Otto, M., Community-associated MRSA: what makes them specialInterestingly, deletion of clfAInterestingly, deletion of clfA Int J Med Microbiol, 2013. 303(6-7): p. 324–30.

13. Lopez, R., et al., Suppression of the lytic and bactericidal effects of cell wallinhibitory antibiotics. Antimicrob Agents Chemother, 1976. 10(4): p. 697–706.

14. Brooks, T. and C.W. Keevil, A simple artificial urine for the growth of urinary pathogens. Lett Appl Microbiol, 1997. 24(3): p. 203–6.

15. Bouatra, S., et al., The human urine metabolome. PLoS One, 2013. 8(9): p. e73076.

16. Stauff, D.L. and E.P. Skaar, The heme sensor system of Staphylococcus aureus. Contrib Microbiol, 2009. 16: p. 120–135.

17. Hammer, N.D. and E.P. Skaar, Molecular mechanisms of Staphylococcus aureus iron acquisition. Annu Rev Microbiol, 2011. 65: p. 129–47.

18. Roos, V., et al., The asymptomatic bacteriuria Escherichia coli strain 83972 outcompetes uropathogenic E. coli strains in human urine. Infect Immun, 2006. 74(1): p. 615–24.

19. Ipe, D.S., et al., Discovery and Characterization of Human-Urine Utilization by Asymptomatic-Bacteriuria-Causing Streptococcus agalactiae. Infect Immun, 2016. 84(1): p. 307–19.

20. Berube, B.J. and J. Bubeck Wardenburg, Staphylococcus aureus alpha-toxin: nearly a century of intrigue. Toxins (Basel), 2013. 5(6): p. 1140–66.

21. von Eiff, C., et al., Prevalence of genes encoding for members of the staphylococcal leukotoxin family among clinical isolates of Staphylococcus aureus. Diagn Microbiol Infect Dis, 2004. 49(3): p. 157–62.

22. Jenkins, A., et al., Differential expression and roles of Staphylococcus aureus virulence determinants during colonization and disease. mBio, 2015. 6(1): p. e02272–14.

23. Supersac, G., et al., Assessment of the role of gamma-toxin in experimental endophthalmitis using a hlg-deficient mutant of Staphylococcus aureus. Microb Pathog, 1998. 24(4): p. 241–51.

24. Malachowa, N., et al., Global changes in Staphylococcus aureus gene expression in human blood. PLoS One, 2011. 6(4): p. e18617.

25. Nilsson, I.M., et al., Alpha-toxin and gamma-toxin jointly promote Staphylococcus aureus virulence in murine septic arthritis. Infect Immun, 1999. 67(3): p. 1045–9.

26. Parker, D. and A. Prince, Immunopathogenesis of Staphylococcus aureus pulmonary infection. Semin Immunopathol, 2012. 34(2): p. 281–97.

27. Ponnuraj, K., et al., A “dock, lock, and latch” structural model for a staphylococcal adhesin binding to fibrinogen. Cell, 2003. 115(2): p. 217–28.

28. Alteri, C.J., S.N. Smith, and H.L. Mobley, Fitness of Escherichia coli during urinary tract infection requires gluconeogenesis and the TCA cycle. PLoS Pathog, 2009. 5(5): p. e1000448.

29. Tenover, F.C. and R.V. Goering, Methicillin-resistant Staphylococcus aureus strain USA300: origin and epidemiology. J Antimicrob Chemother, 2009. 64(3): p. 441–6.

30. Bagale, K., et al., Electronic Cigarette (E-Cigarette) Vapor Exposure Alters the Streptococcus pneumoniae Transcriptome in a Nicotine-Dependent Manner without Affecting Pneumococcal Virulence. Appl Environ Microbiol, 2020. 86(3).

31. Kulkarni, R., et al., Cigarette smoke increases Staphylococcus aureus biofilm formation via oxidative stress. Infect Immun, 2012. 80(11): p. 3804–11.

32. Lather, P., et al., Contribution of Cell Surface Hydrophobicity in the Resistance of Staphylococcus aureus against Antimicrobial Agents. Biochem Res Int, 2016. 2016: p. 1091290.

33. John, P.P., et al., Exposure to moderate glycosuria induces virulence of group B streptococcus. J Infect Dis, 2020.

34. Peignier, A., P.J. Planet, and D. Parker, Differential Induction of Type I and III Interferons by Staphylococcus aureus. Infect Immun, 2020. 88(10).

35. Bolger, A.M., M. Lohse, and B. Usadel, Trimmomatic: a flexible trimmer for Illumina sequence data. Bioinformatics, 2014. 30(15): p. 2114–20.

36. Langmead, B. and S.L. Salzberg, Fast gapped-read alignment with Bowtie 2. Nat Methods, 2012. 9(4): p. 357–9.

37. Liao, Y., G.K. Smyth, and W. Shi, The R package Rsubread is easier, faster, cheaper and better for alignment and quantification of RNA sequencing reads. Nucleic Acids Res, 2019. 47(8): p. e47.

38. Love, M.I., W. Huber, and S. Anders, Moderated estimation of fold change and dispersion for RNA-seq data with DESeq2. Genome Biol, 2014. 15(12): p. 550.

39. Young, M.D., et al., Gene ontology analysis for RNA-seq: accounting for selection bias.Genome Biol, 2010. 11(2): p. R14.

40. UniProt, C., UniProt: a worldwide hub of protein knowledge. Nucleic Acids Res, 2019. 47(D1): p. D506–D515.

41. Tenenbaum, D., KEGGREST: Client-side REST access to KEGG. R package version 1.26.1., 2019.

